# Voltage-Gated Sodium Channel Modulation Differentially Alters ON and OFF Bipolar Cell Contributions to the Rat ERG

**DOI:** 10.64898/2026.03.16.712265

**Authors:** Anuradha Vinayak Pai, Yogesh A. Kulkarni, Jayesh Bellare

## Abstract

**Aim:** The ERG b-wave is primarily attributed to ON bipolar cell activity, while the contribution of the OFF pathway and the differential role of voltage-gated sodium (NaV) channels in these pathways remain unclear. This study investigated whether pharmacological modulation of NaV channels differentially alters ON and OFF cone bipolar cell responses and ERG b-wave amplitudes.

**Methods:** Dark- and light-adapted ERGs were recorded from rats across stimulus intensities spanning rod, mixed rod–cone, and cone pathways (1–1000 lux). ON and OFF cone bipolar cell pathways were pharmacologically isolated using intravitreal cis-PDA. NaV channel activity was modulated via intravitreal administration of lidocaine and lamotrigine (blockers) and veratridine (agonist). Changes in b-wave amplitudes were analysed to assess pathway-specific effects.

**Results:** Both lidocaine and lamotrigine significantly globally reduced ERG b-wave across all stimulus intensities, confirming a role for NaV channels in bipolar cell signalling. Pathway isolation revealed differential effects: lidocaine predominantly suppressed ON pathway, whereas lamotrigine preferentially reduced OFF pathway responses. In contrast, veratridine enhanced both ON and OFF pathway activity. These findings indicate that NaV channel activity in ON and OFF cone bipolar cells can be independently and differentially modulated.

**Conclusion:** The ERG b-wave reflects integrated contributions from both ON and OFF cone bipolar cells. Differential NaV channel modulation alters these pathways distinctly, highlighting sodium channel–mediated mechanisms as potential targets for physiologically relevant retinal stimulation strategies in degenerative retinal conditions.

## 1. Introduction

Hereditary degenerative diseases such as retinitis pigmentosa, age related macular degeneration etc. are associated with progressive loss of photoreceptor cells leading to signal impairment at outer plexiform layer (OPL). These conditions are major causes of blindness worldwide. And important because these diseases are progressive in nature, which means the inner retinal neurons deteriorate slowly and remain comparatively healthy for considerable period (Phillips et al., 2010). Retinal bipolar cells, the first vital element in signal relay therefore are considered good targets for electrical / chemical stimulation in order to restore visual signal transmission.

Researchers worldwide have tried to electrically stimulate retinal bipolar cells of degenerating retina using sub retinal implants but have achieved very limited success. Because retinal bipolar cells encode light signals through opposing cation fluxes in ON and OFF pathways, an implant that could take use of this property might be more effective in restoration of retinal processing since it replicates the normal physiology of these cells The main function of retinal bipolar cells is to process electrochemical signals from pre-synaptic photoreceptors into OFF (Dark) / ON (Light) signals at OPL, collectively represented by b-wave of an electroretinogram (ERG). They are so named because ON bipolar cells (ON CBCs) depolarize while OFF bipolar cells (OFF CBCs) hyperpolarize in response to light. This opposing response to light is because ON CBCs contain mGluR6 receptors which when activated cause influx of unknown cations whereas OFF CBCs contain AMPA/Kainate receptors, which lead to outflow of unknown cations upon activation (Euler and Masland, 2000; Strettoi et al., 2010). Cation flux through these channels is a key step in visual signal transmission across OPL (Pugh & Lamb, 2000). Absence or dysfunctional mGluR6, AMPA, or Kainate receptors compromises synaptic signalling Mutations in mGluR6, for example, selectively disrupt ON CBC transmission and are linked to congenital stationary night blindness.

While the cations contributing to voltage response of retinal bipolar cells is presently unknown, Na^+^ and K^+^ currents have been implicated in rod and certain cone bipolar cells (Karschin and Wässle, 1990; Ma et al., 2003; Ma et al., 2005). Functional NaV channels have been identified in cone bipolar cells of rat, mouse and rabbit retinas (Pan & Hu, 2000; Mojumder et al., 2007). Furthermore, intra-vitreal injection of TTX (tetrodotoxin-voltage gated sodium channel blocker) has been shown to reduce both cone-driven (photopic) (Mojumder et al., 2008) as well as rod driven (scotopic) b-wave (Smith et al., 2013) against different background conditions.

In the present study, we investigated whether ON as well as OFF bipolar cell response to graded light stimuli can be simultaneously and differentially modulated by pharmacological manipulation of NaV channel function. And whether such modulation of bipolar cell response can be achieved across stimulus intensities engaging rod, mixed rod + cone, and cone pathways under different ambient conditions (dark-adapted ERG vs. Light adapted ERG).

To achieve this, ON and OFF cone bipolar cell pathways were isolated by intra-vitreal cis-PDA (cis-2, 3-piperidine-dicarboxylic acid), a glutamate analogue, known to suppress signal transmission between photoreceptors and OFF CBCs, horizontal cells as well as third-order neurons (Xu et al., 2003). We then observed effects of intravitreally injected NaV channel antagonists: Lidocaine, Lamotrigine as well as agonist: Veratridine on ON as well as OFF bipolar response of rat ERG.

Our hypothesis was, if activity of NaV channels on cone bipolar cells can indeed be modulated independent of the receptors (mGluR6 or AMPA/Kainate) to which they are likely to be coupled, then these channels could be stimulated clinically to reintroduce vision. Such information will be useful for researchers interested in stimulating retina through sub-retinal implants in order to restore vision.

## 2. Materials and Methods

### 2.1 Animal experiment

Procedures used in animal experiments were approved by Institutional Animal Ethics Committee of NMIMS [approval number CPCSEA/IAEC/P-16/2019] and conducted following the recommendation of the Association for Research in Vision and Ophthalmology Statement for the Use of Animals in Ophthalmology and Vision Research. Wistar rats (both gender, 2-2.5 months of age and 200-225 gm body weight were housed in a room with 12 hr light and dark cycle. Study animals received food and water ad libitum.

### 2.2 Animal preparation

The recording setup was similar to that previously described for studies in rats (Xu et. al., 2003). Animals were dark-adapted overnight and prepared for recording under red illumination (LED > 620 nm). Scotopic ERGs were performed under dim red illumination for rod ERG (n = 30), in 0.5 to 0.6 log cd/m2 (25-31 lux; mesopic ERG, n = 6) and 0.5 log cd/m2 (316-318 lux; photopic ERG, n = 6) background illumination.

Rats were initially anesthetized with an intraperitoneal injection of ketamine (75-90 mg/kg) and xylazine (10 mg/kg) cocktail under dim red light. Anaesthesia was found to set within 5-10 mins for about 45-60 mins. Since recording session lasted about 2 hrs, anaesthesia was maintained by repeat injection of 50% original dose of ketamine-xylazine every 40 mins via a subcutaneous needle fixed to the flank. Rectal temperature was maintained between 36-37 °C by wrapping the rat in a warm blanket. Pupils were dilated to 5 mm in diameter with topical phenylephrine HCl (2.5%) and tropicamide (0.4%). A drop of lidocaine (0.5%) was used for corneal anaesthesia and to reduce eye movement and irritation. The animal’s head was steady throughout recording sessions.

### 2.3 Procedure for intra-vitreal injection of drugs

Animals were subjected to intra-vitreal injection of drugs prior to ERG recording as per procedure mentioned in Xu et al., 2003. Two percent lidocaine was topically applied to achieve corneal anaesthesia. 100 mM cis-PDA dissolved in 4 µl vehicle (normal saline pH 7.4) volume were injected intra-vitreal using a specially designed pre-fillable cartridge with attached 30G needle which was loaded into a standard insulin pen. Our delivery device ensured precise error free injection volume and could be handled by a single person. One eye was used as control (saline injection) whereas the eye served as test (cis-PDA treated). Concentration of lidocaine injected intravitreally was 85.35 mM, of lamotrigine was 600 µM and veratridine was 3.3 µM Antibiotic (ciprofloxacin 0.3% w/v) was applied at the site of needle insertion with the help of earbud / cotton. Animals were returned to their cage and constantly monitored to rule out eye infection / endophthalmitis due to intravitreal injection procedure. All recordings reported here were made 1–2 h after cis-PDA injection.

### 2.4 Recording system, light source, and calibrations

Rats were placed on a table (provided by SVKM’s NMIMS SPPSPTM, Mumbai) specially designed for acquiring ERG in order to keep head in a steady position and in order to reduce noise originating from respiratory and other movements and hence help reliable recording of ERG signals. Recording electrode was specially fabricated circular gold plated ring with circular opening (4 mm diameter) corresponding to pupil diameter of adult rat eye (4-5 mm when dilated) that allowed for light to pass through and was placed directly over the rat cornea of both eyes. Electrode contact was maintained using artificial tears (0.5% w/v carboxy methyl cellulose). Stainless steel needle electrodes were used as reference as well as ground. Reference electrode was needle inserted into head skin of rat while the ground electrode was needle inserted into skin above rat tail. Recording sessions lasted 4 to 8 h.

### 2.5 Method employed for eliciting ERG / stimulus condition

Protocol for all the ERG experiments were uniform for all groups of animals. ERG was recorded by means of Power Lab Data Acquisition system 2/25 (AD Instruments, New South Wales, Australia) and ERGs were acquired using Lab Chart version 7.3.8 (AD Instruments) for Windows, with a Lab Chart Pro license and sent to the computer for averaging, display, storage, and subsequent analysis. A digital 50-60 Hz notch was applied to eliminate line noise. All the electrodes were connected to a differential amplifier (gain of 10,000; sampling rate set at 10 KHz; band pass filter 0.5 – 2 KHz (bridge amplifier). ERGs were recorded against white flash of 509-515 nm wavelength and light calibrations were performed using a photometer (KUSAM-MECO; KM-LUX-99).

Dark-adapted rat ERG was recorded against single 10 msec (2 Hz) flash of white saturating full-field light stimulus (ranging from 1-1000 lux / −2 to 1 log cds / m^2^; Table 2, 4), similar to the one used and more fully described in Xu et. al., 2003. Stimulus was delivered through custom-designed Ganzfeld stimulator placed parallel to both eyes for simultaneous stimulation. Stimulus strengths, number of flashes as well as flash intervals were controlled through an electrical circuit and verified through photodiode coupled to Power Lab Data Acquisition system. According to Mojumder et. al., 2008, Smith et. al., 2013 and Xu et. al., 2003, the range of scotopic b-wave can be subdivided into three sections in rats according to elicited cellular responses. The range of responses (summarized in Table 1 Supplementary) are as follows: rod driven response, rod cone driven response with less cone contribution and rod cone driven response with more cone contribution. Background luminance’s were kept at dim-red (scotopic), 25-31 lux (mesopic) and 316-318 lux (photopic) steady uniform illumination maintained through specially mounted LEDs over the recording area calibrated with a photometer. Steady backgrounds were present for 15–20 min before accepting ERG recording for analysis (Bui & Fortune, 2006). The intervals between flashes were adjusted so that the response returned to baseline before another stimulus was presented. 50 such successive stimuli were presented to record b-wave amplitudes from normal and cis-PDA treated eye at 2.1 Hz against each stimulus intensity per animal in order to reduce trial-to-trial noise. Test and background stimuli were provided by a Phillips LED colour temperatures of 33008K and 34008K, respectively mounted at pre-determined position above rat head to provide uniform illumination which was measured using digital photometer (KUSAM-MECO; KM-LUX-99).

**Table 1.** Comparison of effects of NaV blockers on ON and OFF cone bipolar response of rat ERG.

| Drug | Effect on NaV Channels | ON CBC Response | OFF CBC Response | Dominant Pathway | Net b-wave |
| --- | --- | --- | --- | --- | --- |
| Lidocaine | Block | ↓↓↓ | ↓ | ON | ↓ |
| Lamotrigine | Stabilize inactivated channels | ~ | ↓ | OFF | ↓ |
| Veratridine | Prolong Opening | ↑↑ | ↑↑↑ | OFF | ↓ (paradoxical) |

### 2.6 Data Collection and Statistics

The b-wave and ON CBC responses were evaluated by measuring response amplitudes against each light intensity. OFF bipolar responses were estimated by subtracting the cis-PDA–isolated ON response from the composite ERG waveform, following the approach described by Xu et al. (2003). Statistical analysis was performed using two-way repeated-measures ANOVA or mixed-effects modelling with Geisser–Greenhouse correction, as appropriate, with individual animals serving as unit of analysis, to assess the effect of light intensity as well as treatment. To directly compare pathway sensitivity to drugs, percent suppression was calculated per animal as 100 X [(control-drug treated)/ control] averaged across light intensities. ON and OFF percent suppression were then compared using paired two tailed t-tests with statistical significance set up at 0.5. All analyses were conducted using Graph Pad Prism. Data are presented as Mean ± SEM.

## 3. Results

### 3.1 Effect of cis-PDA on rat ERG

We injected rat vitreous with 4µl 100mM cis-PDA (cis-2,3-piperidine-dicarboxylic acid), in order to suppress signal transmission from photoreceptors to OFF CBCs (HBCs), horizontal cells as well as between bipolar cells and third order neurons (Xu et al., 2003, Naarendorp et al., 1999). And noted the smooth positive wave responses of the ON CBCs (Slaughter & Miller, 1983) which increased as flash intensity increased and in line with Xu et al., 2003. We could observe best effects of 4µl 100mM cis-PDA between 1.5-3 hrs after injection.

We then proceeded to evaluate effects of NaV channel modulators ON and OFF CBC responses.

### 3.2 Effect of Lidocaine on b-wave of ERG

4µl intra-vitreal Lidocaine (2%) significantly inhibited b-wave of rat ERG (mixed-effects model with Geisser–Greenhouse correction, p= 0.003; Sidak multiple comparisons *post hoc* test 95% CI) (Fig.1). The effect of lidocaine on b-wave was dependent on light intensity (p=0.0489) with significant suppression observed at 130 lux (p=0.0066) despite reductions being seen at all intensities.

**Fig. 1.**
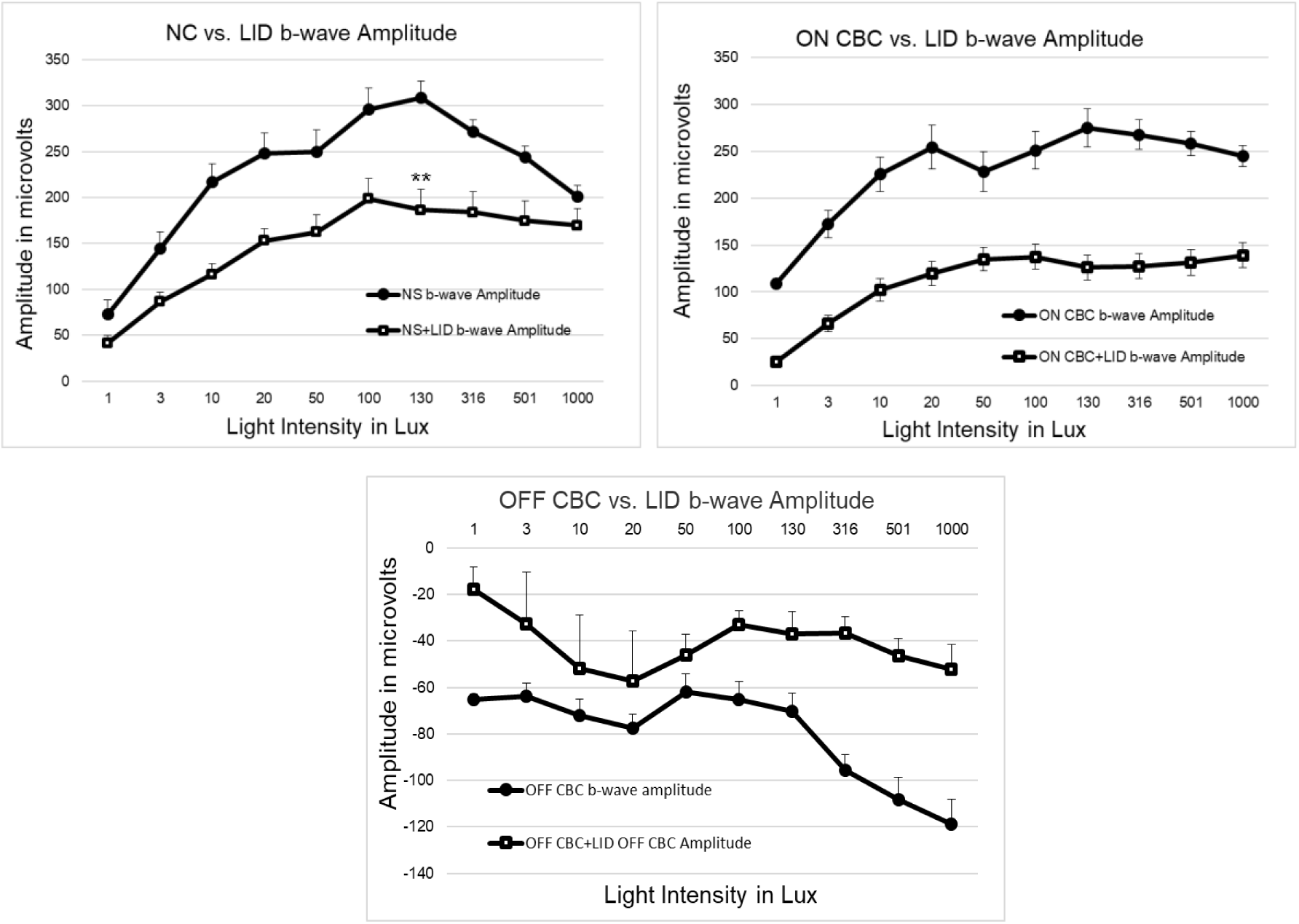
Top Left: Effect of Lidocaine on b-wave of rat ERG against 1, 3, 10, 20, 50, 100, 130, 316, 501 and 1000 lux. ERG recorded simultaneously against white flash stimuli from both eyes of rat; one eye saline-treated control and other eye treated with 4µl intra-vitreal Lidocaine (2%). Top Right: Effect of Lidocaine on ON CBC response. One eye saline+ cis-PDA-treated control and other eye treated with cis-PDA + 4µl intra-vitreal Lidocaine (2%). Bottom: Effect of Lidocaine on OFF CBC response. Lidocaine significantly reduced both ON CBC and OFF CBC activity contributing to reduction in overall b-wave amplitude, corresponding Student’s t-test values have been shown on right.

We then went to observe which pathway (ON vs OFF) accounted for lidocaine-induced reduction in ERG b-wave. And found lidocaine causes a robust and highly significant suppression of ON CBC amplitudes (mixed-effects model with Geisser–Greenhouse correction, p= 0.0002; Sidak multiple comparisons *post hoc* test 95% CI) as compared to OFF CBC amplitudes (p= 0.0264; Sidak multiple comparisons *post hoc* test 95% CI). A percent suppression comparison of ON vs OFF pathway (after normalization) using paired two-tailed student’s t-test (t (8) = 0.64, p = 0.54) however did not show any significant difference.

### 3.3 Effect of Lamotrigine on b-wave of ERG

Similarly, 4µl intra-vitreal Lamotrigine (600µM/ml) also inhibited b-wave of rat ERG (mixed-effects model with Geisser–Greenhouse correction, p= 0.0073; Sidak multiple comparisons *post hoc* test 95% CI) (Fig.2.).

**Fig. 2.**
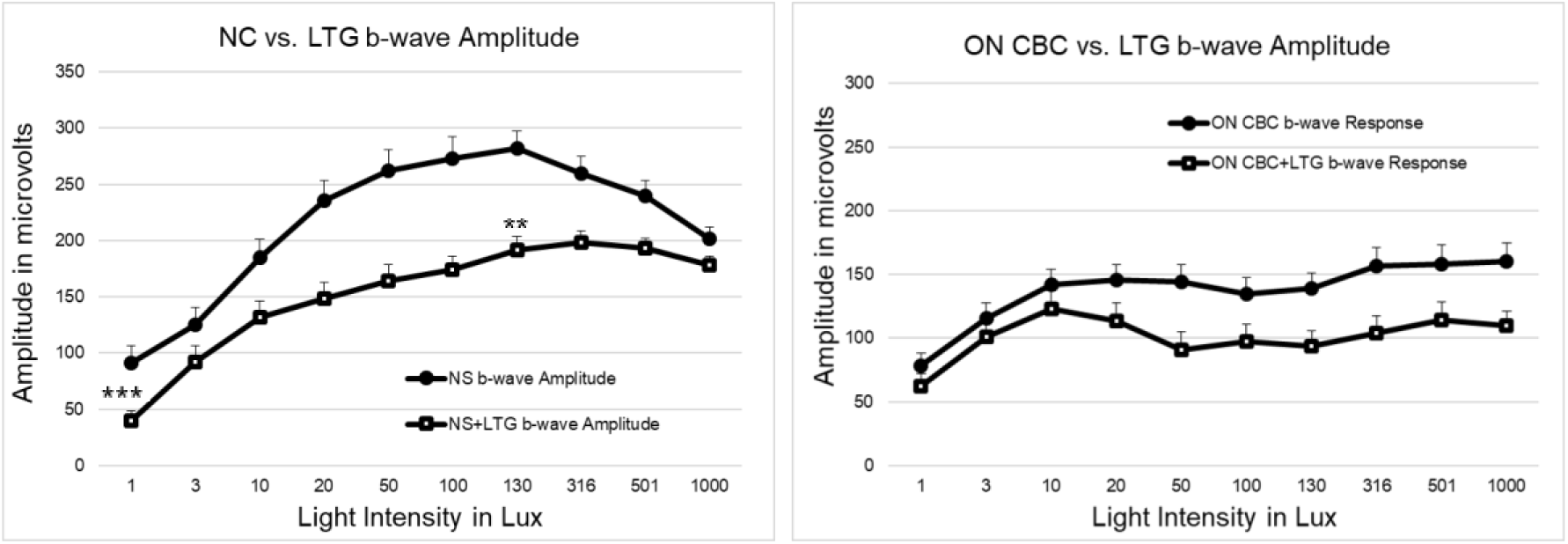
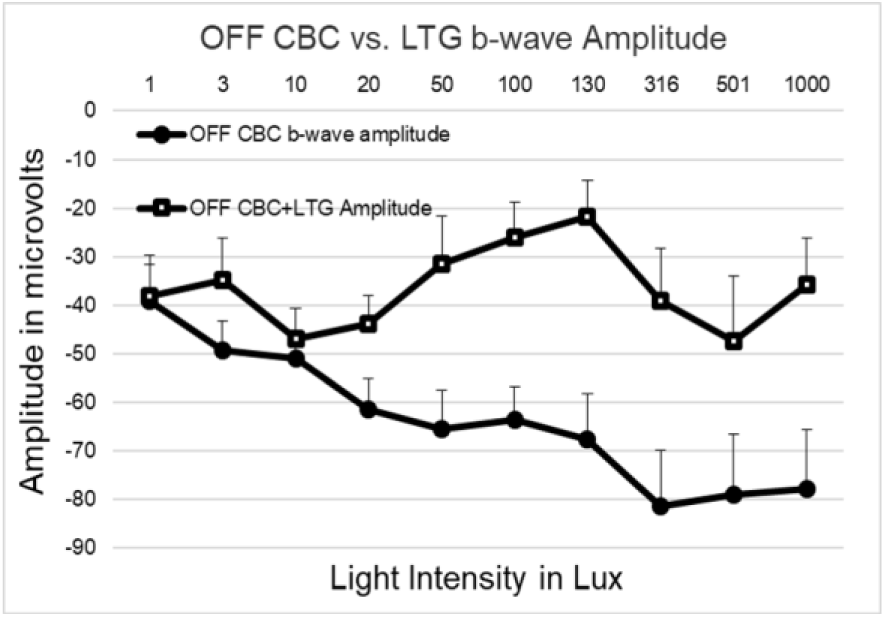
Top Left: Effect of Lamotrigine on b-wave of rat ERG against 1, 3, 10, 20, 50, 100, 130, 316, 501 and 1000 lux. ERG recorded simultaneously against white flash stimuli from both eyes of rat; one eye saline-treated control and other eye treated with 4µl intra-vitreal Lamotrigine (600µM/ml). Top Right: Effect of Lamotrigine on ON CBC response. One eye saline+ cis-PDA-treated control and other eye treated with cis-PDA + 4µl intra-vitreal Lamotrigine (600µM/ml). Bottom: Effect of Lamotrigine on OFF CBC response. Lamotrigine significantly reduced both ON CBC and OFF CBC activity contributing to reduction in overall b-wave amplitude, corresponding Student’s t-test values have been shown on right.

Although the effect of lamotrigine on b-wave amplitudes was moderate as compared to lidocaine, the drug effect was dependent on light intensity (p=0.0113). Significant suppression was observed at 1 lux (p=0.0031) and 130 lux (p=0.0179) despite reductions being directional at all intensities.

We then went to observe which pathway (ON vs OFF) accounted for lamotrigine-induced reduction in ERG b-wave. Although there was numerical reduction of ON CBC responses, the effect was neither significant nor reliable (mixed-effects model with Geisser–Greenhouse correction, p= 0.0589; Sidak multiple comparisons *post hoc* test 95% CI −1.67 to 75.25). However OFF CBC amplitudes were significantly and reliably reduced (p= 0.0224; Sidak multiple comparisons *post hoc* test 95% CI) and the effect was consistent across luminance. A percent suppression comparison of ON vs OFF pathway (after normalization) using paired two-tailed student’s t-test (t (8) = 0.41, p = 0.69) however did not show any significant difference.

Since both sodium channel blockers (lidocaine and lamotrigine seemed to have inhibitory effect on ON and OFF CBCs and therefore on b-wave of rat ERG, we probed whether a sodium channel stimulator would have opposite effect.

### 3.4 Effect of Veratridine on b-wave of ERG

4µl intra-vitreal Veratridine (3.53 mM) surprisingly caused significant luminance dependent suppression of b-wave amplitudes (mixed-effects model with Geisser–Greenhouse correction, p= 0.0330; Sidak multiple comparisons *post hoc* test 95% CI) especially at 500 lux (p= 0.0113) and 1000 lux (p= 0.0223).

However, veratridine stimulated both ON CBC (p= 0.0088) as well as OFF CBC (p= 0.0044) pathway and the effect was consistent across lux (Fig.3.). This was surprising as both ON and OFF CBCs majorly contribute to b-wave of ERG (in the absence of signals from horizontal cells and third-order neurons) and stimulation of both these pathways should have caused enhancement of ERG b-wave.

**Fig. 3.**
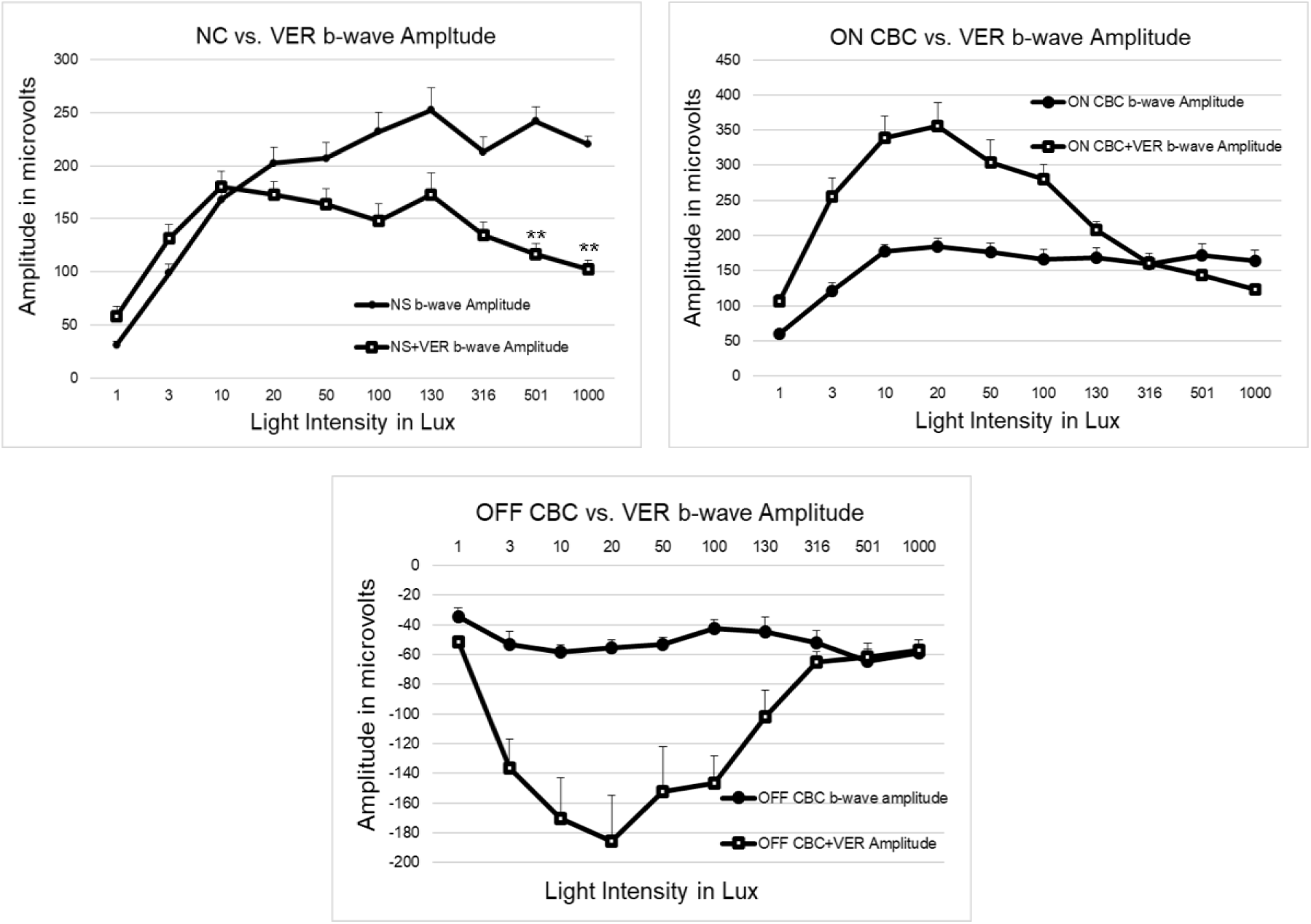
Top Left: Effect of Veratridine on b-wave of rat ERG against 1, 3, 10, 20, 50, 100, 130, 316, 501 and 1000 lux. ERG recorded simultaneously against white flash stimuli from both eyes of rat; one eye saline-treated control and other eye treated with 4µl intra-vitreal Veratridine (3.53 mM). Top Right: Effect of Veratridine on ON CBC response. One eye saline+ cis-PDA-treated control and other eye treated with cis-PDA + 4µl intra-vitreal Veratridine (3.53 mM). Bottom: Effect of Veratridine on OFF CBC response. Although veratridine activates NaV channels and increases amplitudes in both ON and OFF CBCs, preferential stimulation of OFF CBCs lowers the ON: OFF ratio, there is a paradoxical reduction in b-wave amplitude following an initial augmentation.

A percent suppression comparison of ON vs OFF pathway (after normalization) using paired two-tailed student’s t-test (t (8) = 2.53, p = 0.0445) however showed OFF pathway was more significantly affected than ON pathway.

Although the ERG b-wave is classically attributed to ON CBC activity, evidence from lamotrigine and veratridine effects on b-wave indicated that OFF pathway signalling can significantly modulate the net b-wave. Therefore, we compared ON CBC: OFF CBC ratio to assess whether shifts in ON–OFF balance contribute to the observed reduction in b-wave amplitude.

### 3.5 Comparison of ON CBC: OFF CBC ratio between NS, LID, LTG and VER treated retina

We discovered that that ERG b-wave amplitude is critically dependent on the relative balance between ON and OFF pathway activity (Fig.4.). A reduction in the ON: OFF CBC ratio was consistently associated with attenuation of the b-wave. Lidocaine significantly reduced ON CBC activity whereas lamotrigine significantly reduced OFF CBC activity. The net effect however was that b-wave primarily reduced because of reduction in ON: OFF CBC ratio. Whereas a sodium channel activator veratridine stimulated both ON as well as OFF CBCs. But stimulation of OFF CBCs was far more robust counterbalancing ON pathway excitation paradoxically leading to reduced ON: off CBC ratio resulting in overall reduction of ERG b-wave amplitude (Table 1).

**Fig. 4.**
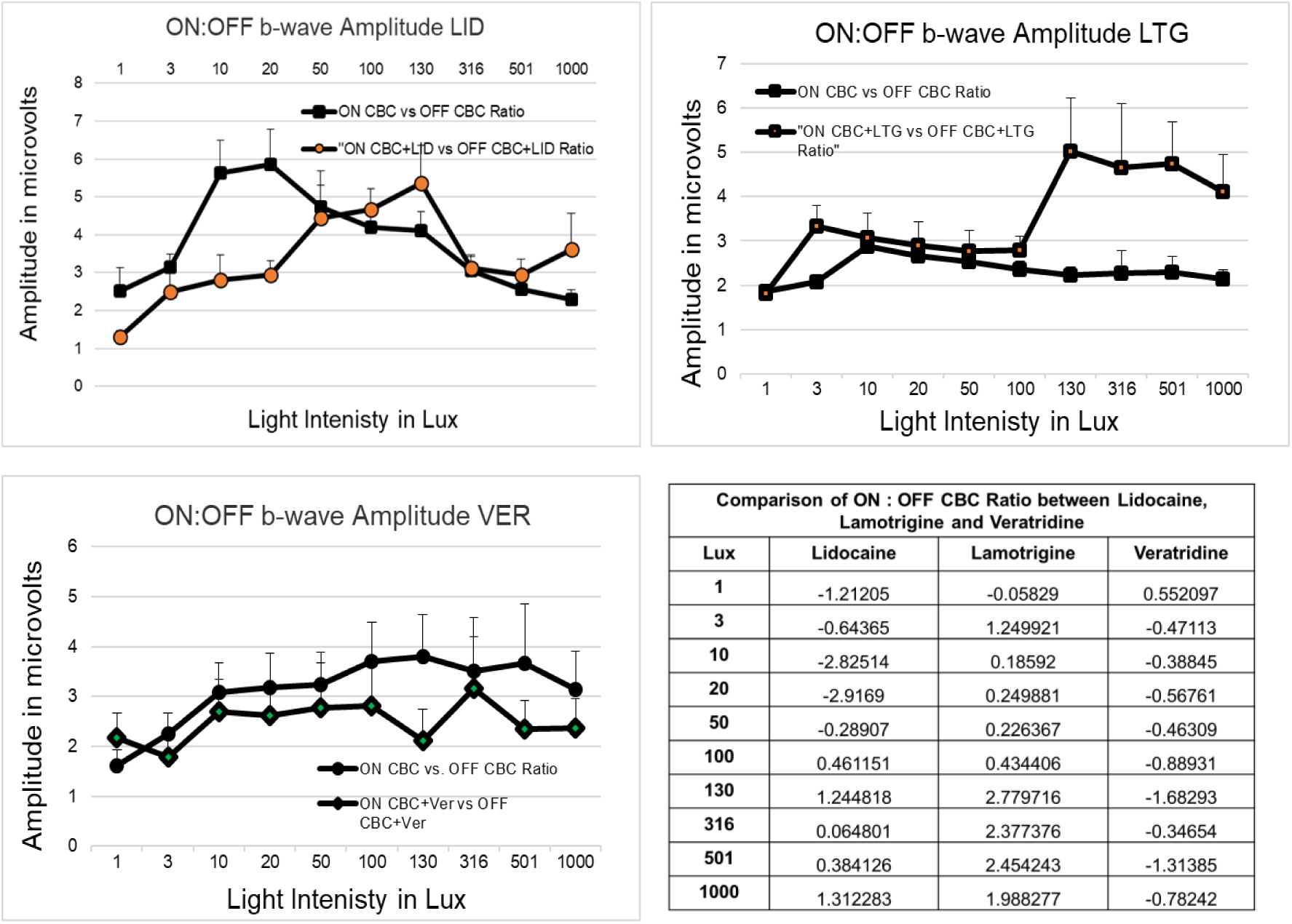
Top Left: Effect of Lidocaine on ON CBC: OFF CBC ratio; Top Right: Effect of Lamotrigine on ON CBC: OFF CBC ratio and Bottom Left: Effect of Veratridine on ON CBC: OFF CBC ratio. A paradoxical reduction in b-wave amplitude following veratridine is due to preferential stimulation of OFF CBCs as compared to ON CBCs. Student’s t-test values have been shown on bottom right.

## 4. Discussion

Light photons strike the photoreceptor detectors of retina and cause reduction in its membrane potential. Photoreceptor hyperpolarization is directly proportional to the amount of photons (light intensity) and directly correlates to reduction in release of glutamate at I plexiform. In retina, a saturated level of photoreceptor glutamate release signals dark and a reduction signals light. Postsynaptic bipolar cells detect changes in this glutamate levels as light or dark. While ON Bipolar cells detect glutamate reduction as light (more reduction = more light; through sign inverting mGluR6 receptors), OFF Bipolar cells detect the same as “reduction in dark” (more reduction = less dark; through sign conserving AMPA/Kainate receptors). Which means the same light signal from photoreceptor cells gets bifurcated into two types of signals: ON bipolar depolarization and reduced OFF bipolar hyperpolarization. The b-wave of an ERG is due to collective response of ON and OFF bipolar cells. Any disease affecting visual function is however confirmed by observing reduction in b-wave of ERG which is inferred to be caused only by ON bipolar cell signalling.

At I plexiform, both ON as well as OFF CBCs respond to single neurotransmitter glutamate and both cells respond by change in cationic flux across cell membrane. Dark causes inflow of unknown cations in OFF CBCs (Puller et al., 2013, Strettoi et al., 2010) whereas light causes inflow of cations in ON CBCs (Bloomfield and Dacheux, 2001; de la Villa et al., 1995). Inflow of cations during depolarization is a universal and fundamental aspect of action potential generation in any neuron. While the cation responsible for depolarization of these bipolar cells is unknown, we assumed that same cation (and perhaps Na^+^ ion) moves across both cells during dark to light activation which governed our choice of using NaV blockers in present study.

Moreover, TTX has been shown to reduce b-wave against different background conditions (Mojumder et al., 2008, Smith et al., 2013). Therefore, we moved forward with the following questions in the current work: Can substances that affect sodium channels be used to modify (inhibit/stimulate) ON bipolar cells? What impact do these substances have on the response of OFF bipolar cells? What impact does the OFF bipolar cell response have on the ERG b-wave? Is it possible to restore bipolar signal transmission by taking advantage of sodium channel function? Our results are as follows:

### 4.1 NaV blockers reduce b-wave response of rat ERG

Both lidocaine and lamotrigine reduced b-wave response of rat ERG. Lidocaine is a local anaesthetic and is known to block NaV 1.1 channels (England & Groot 2009, Scholz et al., 1998, Mojumder et al., 2007, Puthussery et al., 2013) known to be present in I plexiform in either closed or inactivated open state (Matsubara et al., 1987). It is also a known substrate for both TTX sensitive (NaV1.1, NaV1.2, NaV1.3 and NaV1.6) as well as TTX resistant (NaV1.8 and NaV1.9) channels (Strassman & Raymond, 1999) present in various regions of retina and in CNS (Scholz et al., 1998; Brien et al., 2008). Whereas lamotrigine is an anti-epileptic and is known to block pre-synaptic NaV 1.1 channels (Attwell et al., 1998) and is also a known substrate for TTX sensitive NaV1.1, 1.2 and 1.6 channels (Sandalon et al., 2013).

Intraocular lidocaine has been reported to cause reduction in “b” wave amplitudes while intracameral injection has been reported to cause transient loss of visual acuity (Liang et al., 1998, Anders et al., 1999). Similarly, TTX has been shown to suppress rat ERG b-wave by inhibiting NaV channels on ON CBCs (Mojumder et al., 2008). At saturating concentrations, TTX has been additionally shown to reduce b-wave amplitude at stimulus intensities that preferentially activate the rod pathway (Smith et al., 2013).

Our results are consistent with earlier reports. In current investigation, both lidocaine and lamotrigine (also NaV blockers) produced significant reduction of dark-adapted rat ERG b-wave at stimulus strengths ranging from 1 to 1000 lux. These findings indicate sodium channels contribute to bipolar cells signalling across rod, cone as well as mixed rod + cone pathways.

### 4.2 NaV blockers reduce b-wave by reduction of both ON and OFF bipolar cell response

We next examined whether the reduction in the rat ERG b-wave produced by lamotrigine and lidocaine, arises exclusively from suppression of ON bipolar cell activity or also involves inhibition of OFF bipolar cell signalling. To address this, we evaluated the effects of these blockers in retinas treated with cis-PDA.

Lidocaine significantly suppressed both ON and OFF CBC pathways (Fig.5.). However, the magnitude of suppression was substantially greater in the ON pathway, indicating that the drug-induced reduction of the ERG b-wave is mediated predominantly by inhibition of ON CBC signalling. In contrast to lidocaine, lamotrigine did not significantly suppress of ON CBC responses, but produced a significant reduction in OFF pathway activity, indicating that lamotrigine-induced b-wave attenuation is mediated primarily through OFF CBC signalling, with ON responses showing only a non-significant trend (Table 1).

**Fig. 5.**
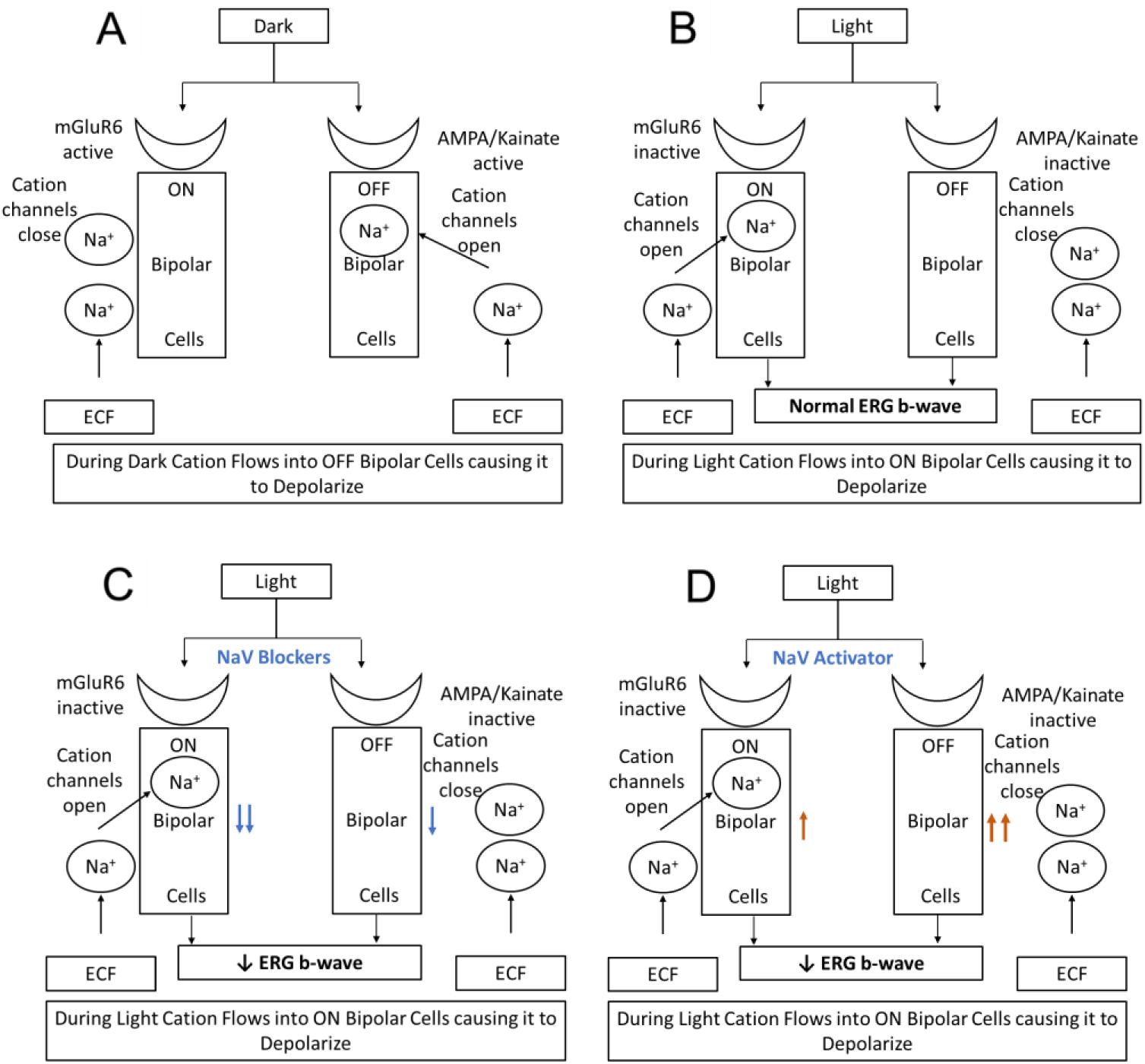
Our view of effect of lidocaine, lamotrigine and veratridine on NaV channels of ON and OFF Cone bipolar cells. A & B: Normal flow of Na^+^ currents in Dark vs. Light. C: NaV blockers Lidocaine & Lamotrigine preferentially suppress ON CBC signalling leading to reduced ERG b-wave. D: NaV activator Veratridine enhances NaV dependent signalling with dominant OFF CBC activation negating ON CBC activation leading to paradoxical reduction in ERG b-wave.

Voltage-gated sodium channels have been reported in retinas of rat, tiger salamander, ground squirrel and rabbit implying significant role of sodium currents in visual processing at I plexiform (Cui & Pan 2008, Becker et.al., 2016). NaV channels have also been identified in both ON and OFF types of CBCs of goldfish retina (Zenisek et al., 2001) and immunolocalized to axon shafts of specific OFF CBCs implicated in dim light visual signalling in mouse retina (Hellmer et. al., 2016). In primates, NaV1.1 channels enhance excitatory signals from bipolar cells to OFF parasol ganglion cells (Puthussery et. al., 2013), while NaV1.6 contribute signal transmission between photoreceptor cells and ON CBCs (Smith & Côté 2012). Together these findings suggest that different NaV channel subtypes may contribute differently to ERG b-wave due to their distinct cellular and subcellular localization, and may account for mechanistically dissimilar effects of both lidocaine and lamotrigine in reducing ERG b-wave.

Our results further suggest NaV channels in both ON and OFF bipolar cells can be differentially suppressed and that suppression of either pathway causes reduction of ERG b-wave amplitudes.

### 4.3 Both ON and OFF bipolar cells can be stimulated by sodium channel opener drug

Intra-vitreal Veratridine stimulated both ON as well as OFF bipolar cells. Veratridine inhibits NaV channel inactivation leading to prolonged channel opening, and is thus employed as neuropharmacological tool to enhance depolarization (Ulbricht W., 1998). It also acts by binding within NaV channels causing a lowering of activation threshold (Craig et. al., 2020) with greater selectivity for TTX-sensitive than TTX-resistant channels (Farrag et. al., 2008), an effect opposite to that of lidocaine and lamotrigine.

In the present study, veratridine increased the response amplitudes of both ON and OFF CBCs. While ON pathway responses were strongly enhanced in a luminance-dependent manner, the magnitude of stimulation was significantly greater in the OFF pathway (Fig.5.). This indicates OFF CBC signalling is the dominant contributor to the veratridine-induced alteration of the ERG b-wave. These results suggest OFF circuitry is also NaV channel dependent and it is possible to independently activate both ON and OFF pathways through stimulation of NaV channels.

### 4.4 Proportional stimulation of ON and OFF bipolar cells is required to restore signal transmission through retinal bipolar cells

Despite stimulation of ON and OFF CBCs, veratridine caused a paradoxical reduction of ERG b-wave amplitude. This was because enhanced stimulation of OFF CBCs overshadowed stimulant effects of ON CBCs. A comparison of ratio of ON CBC vs. OFF CBC indicated that a decrease in the ON: OFF CBC ratio—whether produced by preferential suppression of ON signalling or disproportionate enhancement of OFF signalling—leads to attenuation of the b-wave. Whereas a higher ON: OFF CBC ratio transforms into a larger b-wave amplitude, in conditions that preserve dominant ON CBC responses, as can be seen in case of lidocaine and lamotrigine.

These findings indicate that OFF CBCs are also active contributors to retinal signal integration and function in modulation ERG b-wave magnitude. And that ERG b-wave amplitude is governed by the relative balance of ON and OFF CBC activity rather than by absolute pathway activation alone.

### 4.5 The significance of OFF CBC contribution to ERG

Present study for the first time highlights a previously unappreciated role of OFF CBCs in visual signalling. Earlier it was assumed ON CBCs are principal and sole contributors of ERG whereas OFF CBCs were considered electrically silent contributors to b-wave. In contrast, our findings demonstrate OFF CBCs significantly influence ERG b-wave by modulating, distorting, or counterbalancing ON CBC responses particularly under conditions of altered sodium channel function. Therefore, OFF CBCs are not functionally silent in shaping the ERG b-wave, reiterating an earlier observation that these cells could encode distinct aspects of visual signalling (Hellmer et. al., 2016).

Although the precise cellular mechanisms underlying OFF CBC contributions remain to be defined, our data establish that the ERG b-wave represents a composite response reflecting integrated activity from both ON and OFF CBC pathways.

## 5. Conclusion

Present study establishes sodium (Na⁺) currents are critical contributors to visual signalling within the outer plexiform layer (OPL), and these currents operate with opposite polarity and functional consequences in ON and OFF CBC pathways. Also voltage-gated NaV channels present on bipolar cells (dendrites, soma or axon terminals) not only amplify synaptic transmission at OPL, but also regulate the balance between ON and OFF pathway activity, thereby shaping the final ERG b-wave.

Therefore, the ERG b-wave is not just a simple arithmetic sum of ON + OFF CBC amplitudes but a coordinated balanced activity of both ON and OFF bipolar cells in determining magnitude of ERG b-wave. Thus OFF CBCs play an active role rather than a unidirectional ON pathway output in shaping visual signalling at OPL.

Our results also suggest independent modulation of ON and OFF bipolar cells may be possible through NaV channels, despite the fact that they are tightly coupled to AMPA/Kainate or mGluR6 receptors at ribbon synapses of I plexiform. However, a proportionate stimulation of ON and OFF CBCs through NaV channels is needed and can be attempted to restore ON Bipolar signalling. That could be useful in diseases where there is no pre-synaptic input to these cells due to degeneration of photoreceptors.

## 6. Acknowledgments

This work was supported by a BioCARe grant from the Department of Biotechnology (No. BT/PR19955/BIC/101/589/2016), Ministry of Science & Technology, Government of India. We thank Dr. Yogesh A. Kulkarni (Associate Professor), his student Dr. Alok D. Singh, and Dr. Shailesh Khade (Veterinarian) from Shobhaben Pratapbhai Patel School of Pharmacy & Technology Management (SPPSPTM), SVKM’s NMIMS, Mumbai for their support in helping with animal care and handling. We thank Mr. Shailesh Gala from Ms. Visha World (Lamington Rd, Grant Road East, Mumbai) for providing electronic parts for ERG instrumentation. We thank Dr. Kishore Dave (Shrushrusha Hospital, Dadar, Mumbai) for guidance on electrophysiological measurements. We also thank Mr Chandrashekhar Kulkarni, Associate Professor, Thadomal Shahani Engineering College Mumbai support in electrophysiological setup. for However, the article’s content, views, and conclusions contained herein are those of the primary and corresponding authors and are solely their responsibility.

## 7. Author contributions

Dr. Anuradha V Pai performed the experiments and processed the data. Dr. Jayesh Bellare and Dr. Yogesh A. Kulkarni helped design the experiments. Dr. Jayesh Bellare contributed to data analysis and interpretation. Both Dr. Anuradha V Pai and Dr. Jayesh Bellare wrote the manuscript.

## 9. Additional information

Extended data is available on request from the author.

## 10. Declaration of generative AI and AI-assisted technologies in the manuscript preparation process

During the preparation of this work the author(s) used AI-assisted language tools [Grammarly, Quillbot/ English Grammar Correction] in order to improve grammar and writing clarity. After using this tool/service, the author(s) reviewed and edited the content as needed and take(s) full responsibility for the content of the published article.

## Supplementary

**Table 1.**
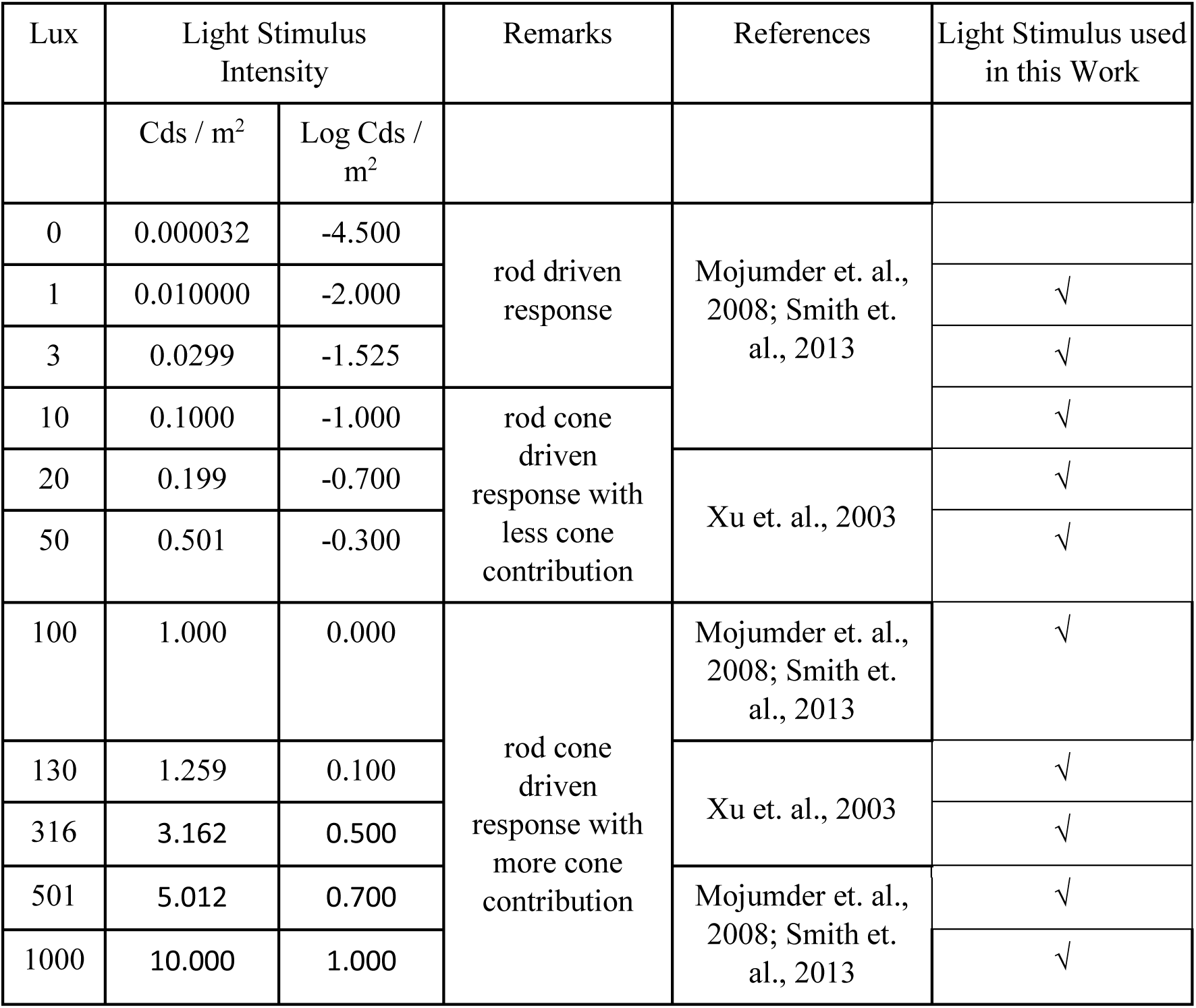
Different stimulus strengths used in the present study to excite rod driven, mixed rod cone driven response with less cone contribution.

